# Identification of Novel Mutations Causing Clofazimine Resistance in *Mycobacterium intracellulare*

**DOI:** 10.1101/2025.04.09.647911

**Authors:** Xiuzhi Jiang, Dan Cao, Xu Dong, Pusheng Xu, Yi Li, Xin Yuan, Yanghui Xiang, Kaijin Xu, Ying Zhang

## Abstract

Clofazimine (CFZ) is a promising repurposed drug for treating *Mycobacterium avium*-*intracellulare* complex pulmonary disease (MAC-PD), but its resistance mechanisms in *M. intracellulare* remain poorly understood. In this study, we generated 36 CFZ-resistant *M. intracellulare* mutants in vitro and identified various mutations in the *marR* gene (WP_009952290.1) in 61% of resistant mutants through whole-genome sequencing. Mutations were identified in additional genes encoding flavin-dependent oxidoreductase (*ssuD*, C67A), membrane lipoprotein (*lppI*, C207 deletion), glucose-methanol-choline oxidoreductase (G157 deletion), MASE1 domain-containing protein (C62G), and PPE family protein (222C deletion). Gene complementation experiments demonstrated that introducing the wild-type *marR* in CFZ-resistant strains (L72 and L74) with *marR* mutations reduced CFZ minimum inhibitory concentrations (MICs) from 1 *μ* g/mL to the susceptible baseline (0.25 *μ* g/mL), confirming its critical role in CFZ resistance. Notably, the *M. intracellulare* MarR lacks homology to *M. tuberculosis* MarR family protein Rv0678 (MmpR) whose mutations are involved in CFZ and bedaquiline (BDQ) resistance and is flanked by non-efflux pump genes (*dhmA* and *doxX*), and unlike *M. tuberculosis*, its mutation does not cause bedaquiline cross-resistance, indicating a different MarR and distinct regulatory mechanism for CFZ resistance in *M. intracellulare*. This work highlights *marR* as a key determinant of CFZ resistance in *M. intracellulare* and underscores the need for further mechanistic studies with implications for clinical diagnostics and therapeutic strategies.

## Introduction

*M. intracellulare* is a member of the *Mycobacterium avium*-*intracellulare* complex (MAC), which belongs to slow-growing non-tuberculous mycobacteria (NTM). It is found in soil, water and animal excreta. Infected birds, mammals, and soil rich in bird droppings may be natural hosts for the bacterium. *M. intracellulare* can cause chronic lung disease (MAC-PD) and poses a serious threat in immunocompromised individuals. Most MAC patients have a five-year all-cause mortality rate of more than 25 percent[1]. According to the 2020 Official ATS/ERS/ESCMID/IDSA Clinical Practice Guideline[2], a three-drug regimen i.e. macrolide plus rifampicin and ethambutol is recommended for patients with MAC lung disease who are susceptible to macrolides, and treatment continued for at least 12 months after negative culture. The recommended three-drug regimen promotes sputum culture conversion but may not reduce the incidence of hospitalization and mortality[3]. However, it has also been shown that patients with cavitary MAC-PD treated with macrolide-containing regimens for ≥ 6 months have lower mortality rates in patients who had sputum culture conversion than patients who did not achieve culture conversion[4]. Regardless, the rate of treatment failure and recurrence of lung disease caused by *M. intracellulare* remains high, and the treatment success rate of the multi-drug regimens containing macrolides is only about 60 percent[5, 6], and new drugs are urgently needed for more effective treatment.

Clofazimine, a phenazine dye with antimicrobial and anti-inflammatory properties, was initially developed in the 1950s for the treatment of leprosy and tuberculosis[7-9]. Subsequent studies revealed its broad-spectrum antimycobacterial activity, coupled with advantages such as a favorable pharmacokinetic profile, minimal dose-dependent side effects, and low cost. In recent years, CFZ has emerged as a promising agent for MAC-PD therapy and is frequently incorporated into alternative regimens. The precise antimicrobial mechanism of CFZ remains incompletely understood, but current research suggests it exerts its effects through multiple pathways. Due to its lipophilic nature, the drug primarily targets the bacterial outer membrane. On one hand, CFZ undergoes redox reactions to generate reactive oxygen species (ROS), inducing oxidative stress[7-9]. On the other hand, it interacts with membrane phospholipids to produce antimicrobial lysophospholipids[9], which synergistically disrupt membrane integrity, interfere with ion transport, and impair ATP synthesis. Collectively, these mechanisms lead to bacterial membrane dysfunction and energy metabolism collapse, ultimately resulting in bactericidal effects. Clinical studies demonstrate that CFZ-containing regimens achieve higher sputum culture conversion rates and therapeutic efficacy in MAC-PD patients ineligible for standard therapies, particularly those with baseline CFZ MIC values ≤ 0.25µg/mL[10]. In severe MAC-PD cases, CFZ exhibited favorable outcomes, with a sputum culture conversion rate of 71% when administered for >6 months[11]. A randomized controlled trial further indicated comparable efficacy and safety profiles between CFZ-ethambutol-macrolide regimen and standard therapy[12]. Although additional clinical studies are needed to validate its long-term efficacy and safety, CFZ represents a critical alternative for MAC-PD management, especially when conventional treatments are contraindicated or ineffective.

An in vitro drug susceptibility study evaluated the MICs of *M. intracellulare* (n=81) to 17 antimicrobial agents. The results demonstrated that the MIC range of clofazimine (CFZ) for susceptible strains was ≤0.25 μg/mL[13]. Currently, the Clinical and Laboratory Standards Institute (CLSI) has not established CFZ breakpoints for MAC, and the mechanisms underlying CFZ resistance in *M. intracellulare* remain poorly characterized. To address this knowledge gap, the present study induced CFZ-resistant *M. intracellulare* mutants in vitro and employed gene complementation test to identify and validate genetic loci linked to CFZ resistance.

## Materials and methods

### Strains, growth conditions, and reagents

In this study, *M. intracellulare* strain 1576, a clinical isolate maintained in our laboratory, was employed for the selection of drug-resistant mutants. This strain was grown in Middlebrook 7H9 broth supplemented with 0.05% Tween 80 and 10% oleic acid, albumin, dextrose, catalase (OADC enrichment) at 37°C. The chemically competent *E. coli* DH5α cells, used for making constructs for genetic complementation experiments, were commercially obtained from Nanjing Vazyme Biotechnology Co., Ltd. The *E. coli-mycobacteria* shuttle plasmid pMV306 to make the *marR* recombinant plasmid pMV306-*marR*, and transformed into *E. coli* DH5 α. The recombinant pMV306-*marR*/DH5 α was grown in LB broth at 37 ° C for plasmid prep for subsequent complementation of the *marR* mutant (see below). All the antibiotics used were purchased from Shanghai Maikelin Biochemical Technology Co.,Ltd. The antibiotics were dissolved in Dimethyl Sulfoxide (DMSO) to prepare a stock solution with an initial stock concentration of 5 mg/mL.

### Drug susceptibility testing

The minimum inhibitory concentrations (MICs) of CFZ against *M. intracellulare* 1576 and its derived mutant strains were determined using the microdilution method (following the CLSI antimicrobial susceptibility testing standard M24-A2) using cation-adjusted Mueller-Hinton broth (CAMHB) supplemented with 5% OADC as the growth medium. CFZ was serially diluted twofold in a 96-well plate, followed by addition of a prepared bacterial suspension to achieve final CFZ concentrations ranging from 8 to 0.0039 μg/mL and a final bacterial concentration of 1×105 to 1×106 CFU/mL. The plates were incubated at 37 °C for up to 14 days, after which bacterial growth at the bottom of the wells was assessed visually. To enhance the visualization of bacterial growth inhibition, 20% Alamar Blue dye was added to each well, followed by incubation for 2–3 hours at 37°C. The color change (from blue to pink) indicates metabolic activity, thereby facilitating the determination of MIC.

### CFZ-resistant mutant isolation

Approximately 2-week-old (stationary phase) cultures of *M. intracellulare* 1576 grown in 7H9 liquid medium were spread onto 7H11 plates (supplemented with 10% OADC) containing 0.25, 0.5, 1, and 2μg/mL of CFZ. After incubating at 37°C for 3 weeks, mutant colonies were picked by transferring them to new plates containing CFZ at the same concentrations as those used for the induction of CFZ-resistant strains to confirm their resistant phenotype. Throughout the process, the wild-type *M. intracellulare* 1576 was retained as a control. It did not grow on the plates containing CFZ. Only those mutant strains that were consistently resistant to CFZ were further analyzed by PCR and DNA sequencing as described below.

### Whole-genome sequencing

Bacterial colonies grown on 7H11 plates containing CFZ were harvested using an inoculating loop and transferred to a 1.5 mL Eppendorf tube containing 500 μ L of TE buffer. The bacterial suspension was then heat-inactivated at 80°C for 30 minutes in a waterbath prior to whole-genome sequencing. For liquid cultures, 2 mL of bacterial suspension with a turbidity of 1 McFarland Standard was centrifuged at high speed, and the supernatant was discarded. The bacterial pellet was resuspended in 500 μL of TE buffer and subjected to heat inactivation at 80 ° C for 30 minutes and sent for whole-genome sequencing at Shenzhen United Biomedical Co., Ltd. To preserve CFZ-resistant mutant strains, 500 μ L of bacterial suspension was mixed with an equal volume of 60% glycerol in cryogenic tubes and stored at -80 °C for future use.

### PCR and DNA sequencing

Colony PCR was performed on the *M. intracellulare* CFZ-resistant mutants isolated in vitro along with sensitive control strain (Con2). The *marR* gene was amplified using the *marR* F (5’ - CATCAACGCCAGCCACTTCATC-3’) and *marR* R (5’-CAGCAGCATGGTGTCGTGACT TC-3’) primers. 2XPhanta Flash Master Mix was used for PCR amplification with parameters as follows: heat denaturation at 98°C for 30 min followed by 30 cycles of 98°C for 10 sec, 55°C for 5 sec, and 72°C for 5 sec and then extension at 72°C for 1 min. PCR amplification products were subjected to Sanger sequencing to confirm the mutations in *marR* gene in selected mutants. Sequencing results were analyzed using SnapGene 6.0.2 software and compared with the sequence of the Con2.

### Complementation of the *marR* gene

Genomic DNA was isolated from the sensitive control strain Con2 using the FastPure® Bacteria DNA Isolation Mini Kit (DC103; Vazyme, China). Gene-specific primers (marRpMV306F: 5′-caggaattcgatatcaagcttCAGGAGGACCGCGGACCC-3′; marRpMV306R: 5 ′-tacgtcgacatcga taagcttTGTCGTGACTTCGCGCCG-3 ′) were designed using the Vazyme Single Fragment Cloning Primer Design Tool to amplify the wild-type *marR* gene for subsequent cloning into the plasmid vector pMV306. The HindIII restriction site within the plasmid was selected as the cloning site to ensure compatibility with the primer-design strategy. PCR amplification of the *marR* gene was performed under the following conditions: initial denaturation at 98°C for 30 s, followed by 35 cycles of denaturation (98°C for 10 s), annealing (60°C for 5 s), and extension (72°C for 4 s), with a final extension step at 72°C for 1 min. Amplified products were resolved by 1% agarose gel electrophoresis in TAE buffer (120 V, 30 min). The target DNA fragment corresponding to *marR* was excised and purified using the FastPure® Gel DNA Extraction Mini Kit (DC301; Vazyme, China). DNA concentration was quantified using a NanoDrop™ One/OneC Spectrophotometer (Thermo).

The plasmid pMV306 was digested with HindIII restriction enzyme (New England Biolabs, USA) at 37 ° C followed by cloning of the *marR* into linearized pMV306 DNA using the ClonExpress® II One Step Cloning Kit (C112; Vazyme, China), in a reaction mixture containing 5.5*μ*L pMV306 DNA, 1.53μL marR DNA, 4μL 5× CE II Buffer, 2μL Exnase II, and nuclease-free water (to a total volume of 20μL). The reaction was incubated at 37°C for 30 min and then cooled on ice or at 4°C. For transformation, 10μL of the recombinant product was added to 100 μL of thawed DH5α competent cells on ice and incubated for 30 min. Cells were DMSO-shocked at 42°C for 45 sec in a water bath and immediately placed on ice for 2-3 min. Subsequently, 900 μ L of antibiotic-free LB broth was added, and the cells were incubated at 37°C with shaking (200-250 rpm) for 1 h followed by plating on LB agar plates containing kanamycin (100μg/mL) and incubated at 37°C for 12–16 h. For the identification of recombinant products, we used SnapGene software to design upstream and downstream primers (marRpMV306FT: 5 ′ -gcttcttgcactcggcatag-3 ′ ; marRpMV306RT: 5 ′ -gatgcctggcagtcgatc-3 ′). The sequences between the upstream and downstream primers encompassed both the plasmid sequence and the *marR* gene sequence. Single colonies were picked from the transformation plates of the recombinant reaction for colony PCR and Sanger sequencing to verify the correct plasmid construct containing the *marR* gene. Then, the recombinant plasmid (named *marR*-pMV306 plasmid) was transformed into CFZ-Resistant mutants L72 and L74 by electroporation as described[14]. Following electroporation, single colonies were picked from the complementation plates of L72, L74, and the vector plasmid control and subjected to colony PCR using primers marRpMV306FT and marRpMV306RT. The PCR products were sent for Sanger sequencing for verification. Subsequently, similar to the MIC determination described above, the MIC levels of the *marR* gene complementation strains, drug-resistant strains L72 and L74, sensitive strain Con2, and the vector plasmid control strain K1 were determined. Additionally, Alamar Blue dye was used for staining to facilitate visual observation.

## Results

### Clofazimine MIC for *M. intracellulare* and isolation of CFZ-resistant mutants

The MIC of *M. intracellulare* 1576 to CFZ was determined using the methodology outlined in the Methods section, revealing an MIC of 0.25 µg/mL. Subsequent CFU enumeration indicated that approximately 5 × 10^7^ *M. intracellulare* bacteria were inoculated onto agar plates containing varying concentrations of CFZ for the purpose of mutant isolation. Following a 3-week incubation period, 36 CFZ-resistant mutants were isolated from plates containing 1 µg/mL CFZ, whereas no growth was observed on plates containing 2 µg/mL CFZ. The mutation frequency conferring resistance to 1 µg/mL CFZ was calculated to be 7×10^−7^. The MIC values of these resistant mutants, along with the susceptible control strain Con2, were further assessed using the microdilution method. The results demonstrated that the MIC of the resistant mutants (1 µg/mL) was fourfold higher than that of the sensitive control strain Con2 (0.25 µg/mL).

### Whole-genome sequencing identified novel genes associated with CFZ resistance in *M. intracellulare*

To identify possible new mechanisms of CFZ resistance, we subjected the 36 resistant mutants to whole-genome sequencing. Genome sequence comparison of the resistant mutants and parental strain Con2 found that various mutations were present within the *marR* gene (NC_016947.1 2172849-2173334 (-)(486 bp), encoding a 161 amino acid protein (WP_009952290.1) in 22 of the 36 CFZ-resistant mutants. These mutations included promoter deletions, premature stop codons, and frameshift mutations (Table 1). In the *marR* gene, the most prevalent mutation identified was duplication of the adenine (A) base at position 91, which was detected in 5 out of the 36 mutant strains. This mutation resulted in a frameshift mutation initiating from serine at position 31. The second most frequent mutation involved the duplication of the cytosine (C) base at position 184, observed in 3 out of the 36 strains, causing a frameshift mutation starting from glutamine at position 62. All other mutation types were each present in only a single strain. Although the frequency of each individual mutation form was relatively low, the cumulative frequency of all mutation forms reached 61% (Table 1). Interestingly, we found that the *marR* gene in our study exhibits no homology with Rv0678/MmpR involved in CFZ resistance in *M. tuberculosis*, and that unlike *M. tuberculosis* where efflux genes are located adjacent to *Rv0678/mmpR*, no efflux genes were found nearby the *M. intracelluare marR* gene (Figure 1).

**Table 1.**
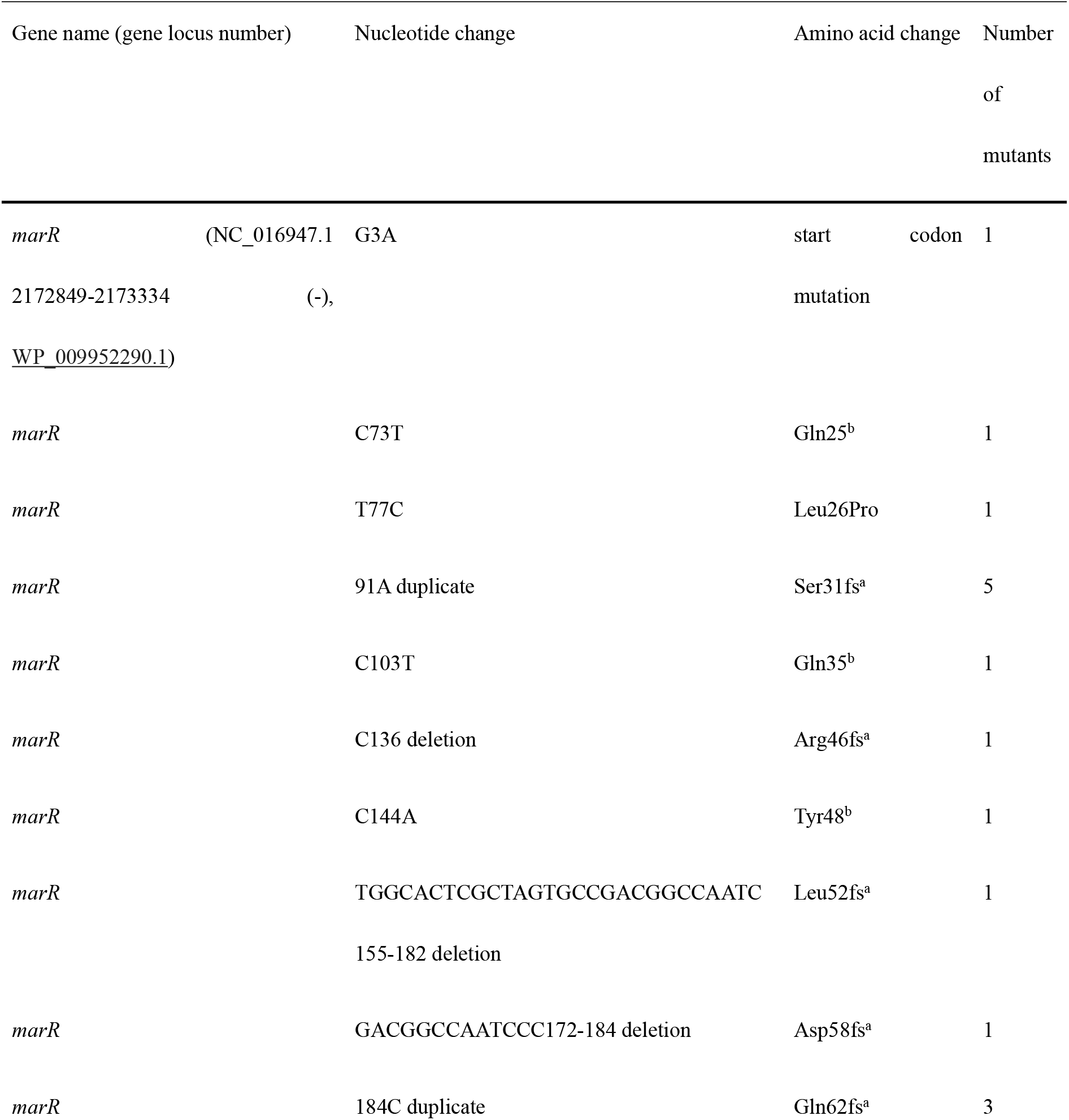
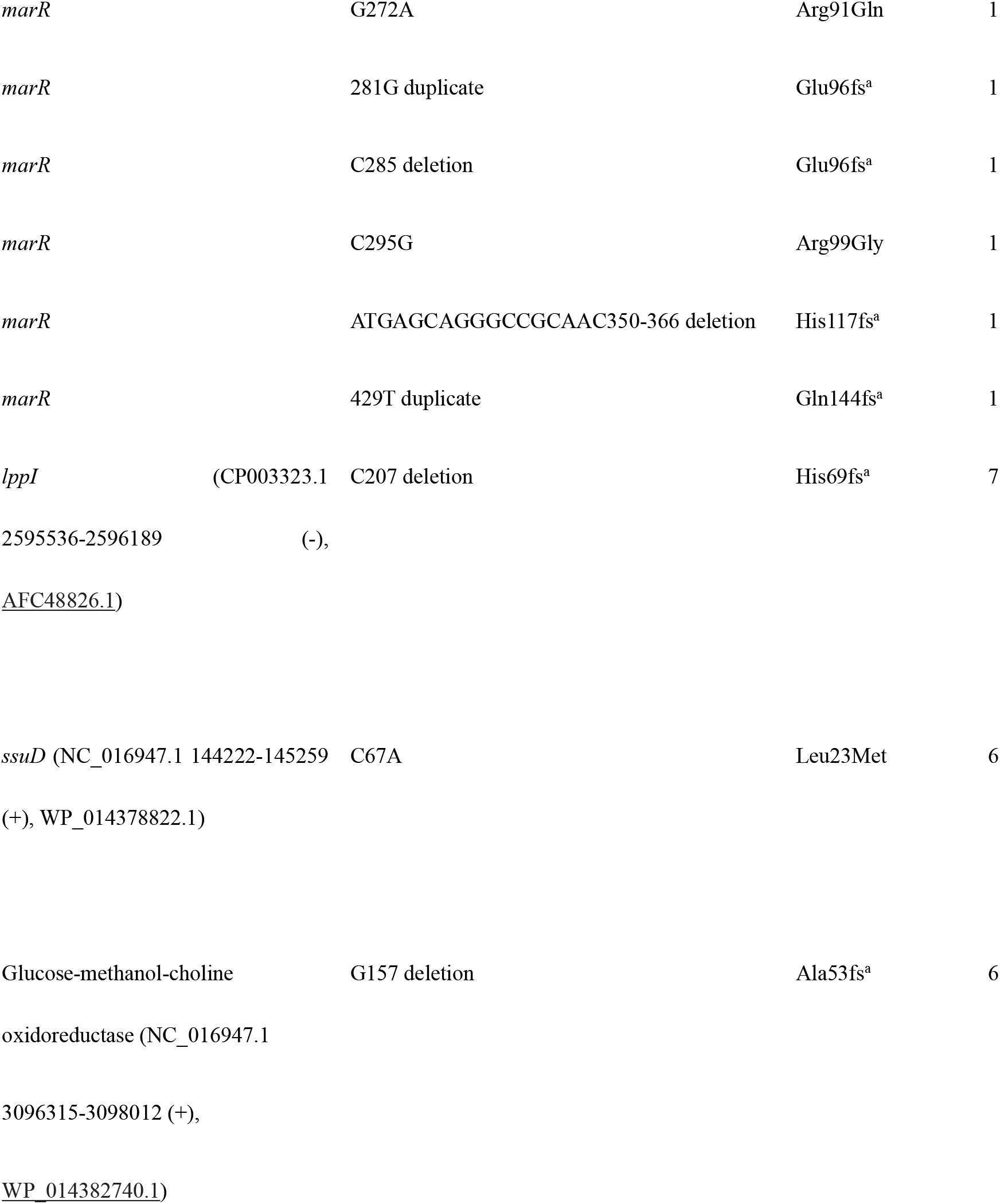
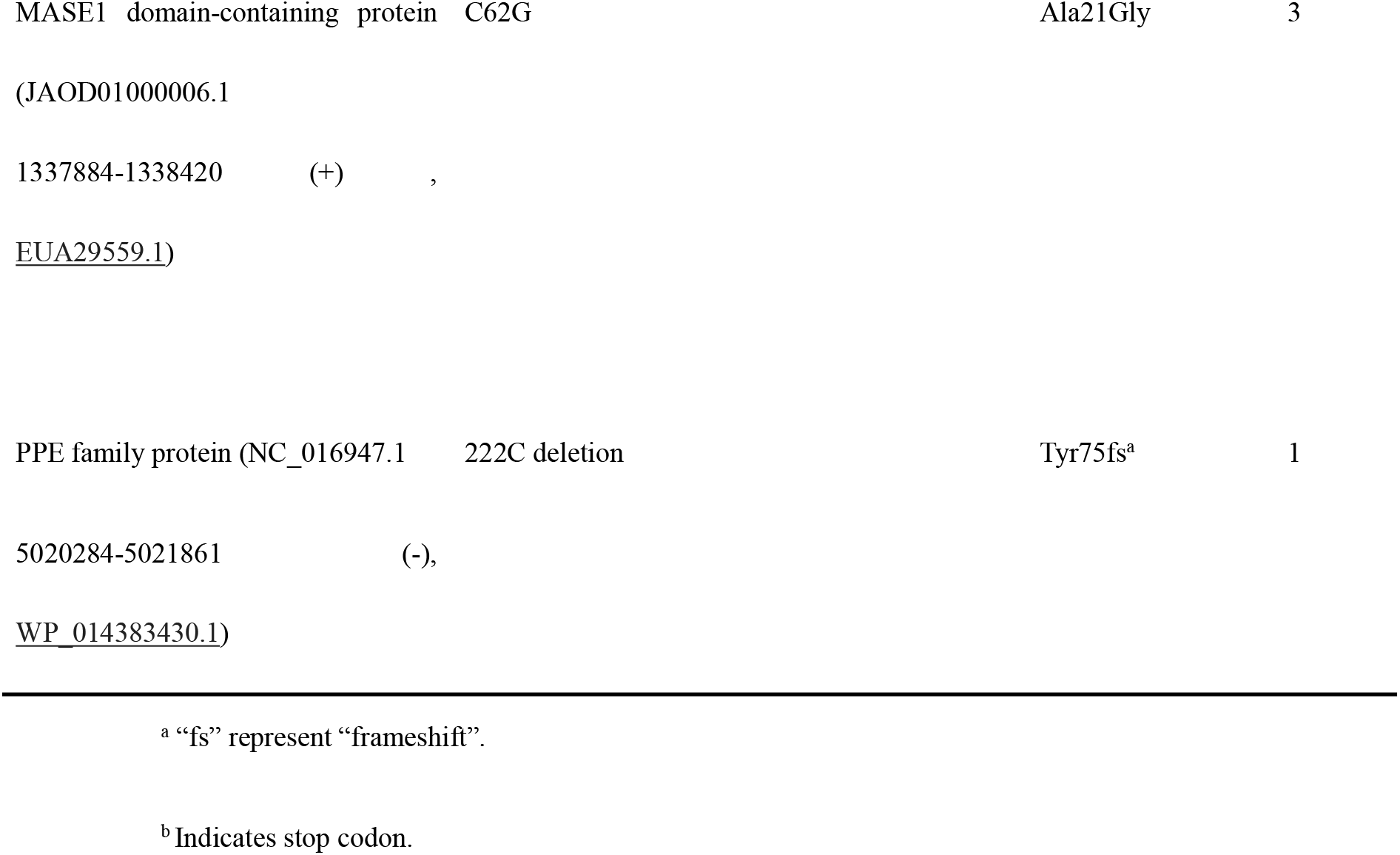
Mutations identified from 36 CFZ-resistant *M. intracellulare* mutants by whole-genome sequencing.

**Figure 1.**
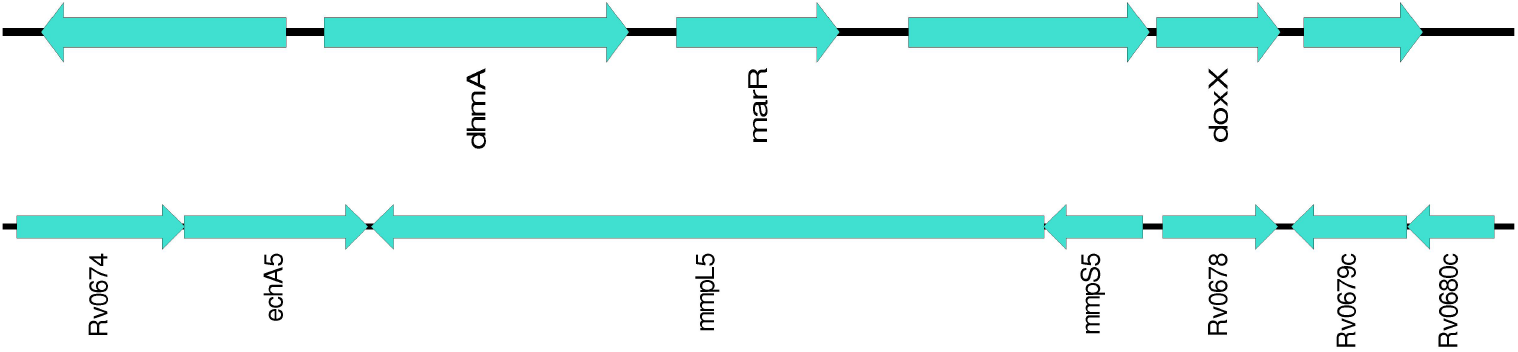
The genome structure diagram of *marR* in *M. intracellulare* 1576 and Rv0678 in *M. tuberculosis*. In *M. tuberculosis* (bottom panel), mutations in Rv0678 (*mmpR/mmpR5*) cause overexpression of its upstream efflux pump genes *mmpS5-mmpL5*, thereby conferring resistance to CFZ. In contrast, the *marR* gene in *M. intracellulare* (top panel) lacks adjacent efflux pump genes. Its upstream gene, *dhmA*, encodes a haloalkane dehalogenase, while the downstream gene, *doxX*, encodes an integral membrane protein.

We also identified that among the 36 mutant strains, 7 strains harbored mutations in the *lppI* gene (CP003323.1 2595536-2596189 (-), AFC48826.1), which encodes lipoprotein LppI. These mutations involved a deletion of the C base at position 207, resulting in a frameshift mutation starting from histidine at position 69. Additionally, 6 out of the 36 mutant strains exhibited mutations in the *ssuD* gene (NC_016947.1 144222-145259 (+), WP_014378822.1), which encodes a flavin-dependent oxidoreductase (Table 1). Specifically, a C-to-A substitution at position 67 led to the replacement of leucine at position 23 with methionine. Among the 7 mutant strains with *lppI* gene mutations, 5 also harbored mutations in the *marR* gene. Similarly, out of the 6 strains carrying *ssuD* gene mutations, 5 exhibited concurrent *marR* mutations. Furthermore, a deletion of the G base at position 157 in the gene encoding glucose-methanol-choline oxidoreductase (NC_016947.1 3096315-3098012 (+), WP_014382740.1) caused a frameshift mutation starting from alanine at position 53. A C-to-G substitution at position 62 in the gene encoding a MASE1 domain-containing protein (JAOD01000006.1 1337884-1338420 (+), EUA29559.1) resulted in the replacement of alanine at position 21 with glycine. Lastly, a deletion of the C base at position 222 in the gene encoding a PPE family protein (NC_016947.1 5020284-5021861 (-), WP_014383430.1) induced a frameshift mutation starting from tyrosine at position 75 (Table 1). Additionally, a mutation site encoding the gene product OB-aCoA-assoc domain-containing protein exhibited a mutation frequency of 100%. However, this mutation was identified as a synonymous variant and was subsequently proven to be unrelated to CFZ resistance through genetic complementation experiments. For subsequent genetic complementation experiments, two resistant strains, L72 (C144A, Tyr48 stop codon) and L74 (G272A, Arg91Gln), both carrying mutations in the *marR* gene, were selected. PCR amplification was conducted on L72, L74 and the control strain Con2 and Sanger sequencing of the PCR products confirmed the accuracy of the mutations.

### *marR* gene complementation and phenotypic validation of the complemented strains

Four single colonies were selected from the *marR* complementation plates of the drug-resistant strains L72 and L74, respectively, and subjected to colony PCR and Sanger sequencing. The sequencing-confirmed complemented strains were designated as 72b, 72c, 74a, 74b, 74c, and 74d. Strains 72b, 72c, 74a, 74b, and 74d were chosen for subsequent phenotypic validation. The minimum inhibitory concentrations (MICs) of these complemented strains, the parental drug-resistant strains (L72, L74), the susceptible control strain Con2, and the empty vector control (K1) were determined using the microdilution method combined with Alamar Blue staining, as described in Methods section. The MIC values of the complemented strains (72b, 72c, 74a, 74b, 74d) decreased to 0.25µg/mL CFZ, equivalent to that of the susceptible strain Con2. In contrast, the MICs of the drug-resistant strains (L72, L74) and the empty vector control strain (K1) remained unchanged at 1µg/mL CFZ (Table 2). To assess potential cross-resistance to BDQ mediated by *marR* mutations identified in this study, we performed BDQ susceptibility testing on the *marR* mutant strains. The results showed that all CFZ-resistant isolates remained susceptible to BDQ with MICs ≤0.0078 µg/mL (Table 2), indicating that the *marR* mutations do not confer cross-resistance to BDQ in *M. intracellulare*.

**Table 2.** CFZ susceptibility results of the *marR* complemented CFZ-resistant mutant strains.

| Strains | CFZ MIC | BDQ MIC |
| --- | --- | --- |
| Con2 | 0.25 mg/mL | $\leq 0.0078$ mg/mL |
| L72 | 1 mg/mL | $\leq 0.0078$ mg/mL |
| L74 | 1 mg/mL | $\leq 0.0078$ mg/mL |
| 72b | 0.25 mg/mL | $\leq 0.0078$ mg/mL |
| 72c | 0.25 mg/mL | $\leq 0.0078$ mg/mL |
| 74a | 0.25 mg/mL | $\leq 0.0078$ mg/mL |
| 74b | 0.25 mg/mL | $\leq 0.0078$ mg/mL |
| 74d | 0.25 mg/mL | $\leq 0.0078$ mg/mL |
| K1 | 1 mg/mL | $\leq 0.0078$ mg/mL |

The strain designations in table 2 are defined as: Con2 (drug-sensitive control), L72 and L74 (CFZ-resistant mutants), 72b/c and 74a/b/d (pMV306-*marR* complemented strains), and K1 (empty pMV306 vector control).

## Discussion

In this study, CFZ-resistant mutants of *M. intracellulare* were isolated in vitro, and whole-genome sequencing was performed to identify resistance-associated mutations. Sequencing revealed diverse mutation types in the *marR* gene among 36 CFZ-resistant mutants, including nonsense, missense, and frameshift mutations, accounting for 61% of the resistant mutants. Two representative mutants, L72 (C144A, stop codon mutation) and L74 (G272A, Arg91Gln missense mutation), were selected for *marR* complementation experiments. Introducing the wild-type *marR* gene in these resistant strains (MIC=1 µg/mL) reduced their CFZ MICs to the level of the susceptible control strain Con2 (MIC=0.25 µg/mL), confirming the role of *marR* mutations in causing CFZ resistance.

In *M. tuberculosis*, CFZ targets the bacterial electron transport chain, with mechanisms of action involving disruption of the redox cycle through enzymatic reduction of CFZ by NDH-2 (NADH dehydrogenase 2), generation of bactericidal reactive oxygen species (ROS), and induction of membrane instability and dysfunction[15]. CFZ resistance mechanisms in *M. tuberculosis* primarily involve mutations in the transcriptional repressor Rv0678, leading to overexpression of the MmpS5-MmpL5 efflux pump. Rv1979c and Rv2535c (*pepQ*) are also involved in CFZ resistance mechanisms in *M. tuberculosis*.[16]. Additionally, a study has demonstrated that Rv1453 is also associated with CFZ resistance in *M. tuberculosis*[17]. However, in *M. avium* there was no association between CFZ resistance and mutations in the Rv0678 homolog, though 2 highly CFZ-resistant *M. intracellulare* strains (MIC = 8 µg/mL) harbored mutations (Asp92Glu and Ala153Pro) in the Rv0678 homolog[18]. However, the functional correlation between these mutations and CFZ resistance was not experimentally validated. It is worth noting that the *marR* gene involved in CFZ resistance in *M. intracellulare* in our study has no homology with *M. tuberculosis* Rv0678 (*mmpR*). To validate whether *marR* gene mutations in this study would confer cross-resistance to BDQ, we determined the MICs of BDQ against *marR* mutant strains. Unlike *M. tuberculosis* MarR family protein Rv0678 whose mutations cause CFZ and BDQ cross-resistance, all tested CFZ-resistant strains exhibited BDQ MIC values ≤0.0078 µg/mL (Table 2), demonstrating no cross-resistance to BDQ in *M. intracellulare*. Furthermore, no efflux pump-associated genes were identified upstream or downstream of the *marR* gene in *M. intracellulare* (Figure 1). The upstream gene *dhmA* encodes a haloalkane dehalogenase involved in detoxifying halogenated hydrocarbons, whereas the downstream gene *doxX* encodes a membrane protein forming a redox enzyme complex with SodA and SseA to mitigate oxidative stress. Although no efflux pump genes flank the *marR* gene, we hypothesize that *marR* mutations in *M. intracellulare* may induce CFZ resistance via upregulation of efflux pump genes located eleswhere in the genome, akin to Rv0678/*mmpR* mutations in *M. tuberculosis*. Transcriptomic studies are warranted to elucidate the regulatory network.

Besides *marR* mutations, we found mutations in *ssuD* gene encoding a canonical flavin-dependent oxidoreductase involved in bacterial alkylsulfonate metabolism to be associated with CFZ resistance in *M. intracellulare*. This enzyme catalyzes the oxidation of alkylsulfonates, enabling bacteria to utilize these compounds as sulfur sources under sulfur-limited conditions. SsuD is indispensable for bacterial metabolism, facilitating both energy production and sulfur assimilation in nutrient-deprived environments. To date, only the *ssuD* gene in Paracoccus sp. strain EF6T has been linked to selenite resistance[19], with no more reported associations between *ssuD* and antimicrobial drug resistance. Another gene involved in CFZ resistance is *IppI* gene which encodes a membrane-anchored lipoprotein critical for cell wall synthesis and membrane integrity in bacteria. LppI contributes to cell wall stability and morphological maintenance by interacting with peptidoglycan layers and membrane-associated proteins. However, no studies have implicated *IppI* in bacterial resistance mechanisms. Glucose-methanol-choline oxidoreductase (GMC oxidoreductase) is a family of redox-active enzymes. Notably, an actinobacteria-independent GMC family oxidoreductase, ChlOR, has been reported to confer resistance to dapsone-like antibiotics. MASE1 domain-containing proteins constitute a class of proteins featuring the “Membrane-Associated Sensor and Effector 1” (MASE1) domain[20]. The MASE1 domain has been identified as a functionally uncharacterized transmembrane domain that typically associates with histidine kinases, GGDEF, GGDEF-EAL, and PAS domains, serving as a sensor to regulate the activity of various output domains[21]. PE/PPE family proteins are critically involved in *M. tuberculosis* pathogenicity. Studies have demonstrated that resistance to the antitubercular compound 3,3-bis-di (methanesulfonyl) propionamide (3bMP1) is conferred by mutations within PPE51, a member of the proline-proline-glutamate (PPE) protein family[22]. Further investigations are required to determine whether the aforementioned genetic mutations contribute to CFZ resistance in *M. intracellulare* in the future.

Although this study has demonstrated that mutations in the *marR* gene of *M. intracellulare* confer resistance to CFZ, several limitations remain. First, this work is primarily based on in vitro-induced drug-resistant mutants, which simulate clinical resistance phenotypes but may exhibit mechanistic differences from drug resistance in authentic clinical isolates. Second, functional validation by complementation was performed only for *marR*, leaving unaddressed the roles of other potential resistance determinants such as *ssuD* and *IppI*. Future investigations should expand the screening of drug-resistant isolates or identify and validate additional resistance-associated genetic alterations from clinical resistant strains. Furthermore, advanced functional assays and mechanistic studies will facilitate a comprehensive understanding of *M. intracellulare* resistance mechanisms, providing critical insights for the development of rapid molecular diagnostics for detecting drug-resistant bacteria.

## Conclusion

This study identified a new mechanism of CFZ resistance mediated by *marR* mutations as a primary driver of resistance in *M. intracellulare*. Through in vitro mutagenesis and functional complementation, we demonstrated that *marR* gene disruption elevates CFZ MICs, while reintroducing the wild-type *marR* gene restores susceptibility. Unlike *M. tuberculosis*, where Rv0678 mutations drive efflux pump-mediated resistance, *marR* in *M. intracellulare* has no homology to *M. tuberculosis* Rv0678 involved in CFZ resistance and is embedded in a genomic context devoid of efflux pump homologs, implying a different resistance mechanism. While mutations in *ssuD, lppI*, glucose-methanol-choline oxidoreductase, MASE1 domain-containing protein and PPE family protein were observed in CFZ-resistant *M. intracellulare*, their roles in CFZ resistance require further investigation. Future studies should prioritize clinical isolates, transcriptomic profiling, and functional validation of different candidate genes in CFZ resistance. Our findings provide critical insights into CFZ resistance mechanism of *M. intracellulare* and have implications for developing rapid molecular susceptibility tests for drug-resistant organisms.

## CRediT authorship contribution statement

**Xiuzhi Jiang:** Conceptualization, Writing – original draft. **Dan Cao:** Supervision. **Xu Dong:** Visualization, Data curation. **Pusheng Xu**: Supervision, Methodology. **Yi Li**: Supervision, Methodology. **Xin Yuan:** Visualization. **Yanghui Xiang:** Visualization. **Kaijin Xu:** Supervision. **Ying Zhang:** Conceptualization, Writing – original draft, Writing – review & editing, Supervision, Funding acquisition.

## Declarations

### Ethical Approval

Not applicable.

### Sequence Information

Not applicable.

### Data availability

Data will be made available on request.

## Acknowledgements

We acknowledge the experimental platform and funding support from Zhejiang University School of Medicine.

